# THE HAPLOTYPE-RESOLVED AND CHROMOSOME-SCALE GENOME OF *VACCINIUM* STAMINEUM: A NEW SOURCE OF GENETIC VARIABILITY FOR BLUEBERRY BREEDING

**DOI:** 10.1101/2025.03.21.644558

**Authors:** Gabriel O. Matsumoto, Juliana Benevenuto, Patricio R. Munoz

**Affiliations:** Blueberry Breeding and Genomics Lab, Horticultural Sciences Department, University of Florida, Gainesville, FL 32611

## Abstract

The *Vaccinium* genus comprises several commercially important fruit crops, such as blueberry, lingonberry, bilberry, and cranberry. However, past breeding efforts have primarily focused on a limited number of wild relatives as sources of genetic variability, leaving a vast genetic reservoir untapped. In this study, we present the first haplotype-phased reference genome for *V. stamineum*, a blueberry wild relative with potential for de novo domestication and introgression into breeding programs. *V. stamineum* is particularly notable for several agronomic traits of interest, such as high soluble sugars, unique flavor profile, and its unique anthocyanin accumulation in fruit pulp, a trait absent in cultivated blueberries.Our assemblies revealed 12 pseudomolecules corresponding to the base chromosome number of *Vaccinium* species, with a genome size of 529.16 Mb and 493.82 Mb for primary and secondary haplotypes, respectively. Despite a slightly smaller genome than other *Vaccinium* species, *V. stamineum* exhibited a higher number of predicted protein-coding genes, while the repetitive elements comprised 39.77% and 42.38% of the primary and secondary haplotypes, respectively. BUSCO analysis indicated 97% transcriptome completeness, supporting the accuracy of gene annotation. Genome-wide alignments showed that *V. stamineum* haplotypes were highly collinear to each other, as well as to *V. corymbosum*. However, further validation is required to resolve a putative translocation in chromosome 1 of the primary haplotype. Altogether, this study established a genomic framework that will facilitate the introgression of traits of interest into blueberry breeding programs and support the potential domestication of *V. stamineum* as a novel fruit crop.

## Background

The *Vaccinium* genus is very diverse, including some important commercial small fruits, such as highbush blueberry (*V. corymbosum* L.), cranberry (*V. macrocarpon* Aiton), bilberry (*V. myrtillus* L.), and lingonberry (*V. vitis-idaea* L.) (Edger et al., 2022). In recent decades, there has been a growing trend towards the consumption of *Vaccinium* berries due to their high levels of health-promoting compounds (IBO, 2023). Among the cultivated species within this genus, highbush blueberry has the most economic importance, though this is a relatively young crop. The domestication of highbush blueberries occurred primarily during the 20th century (Lyrene and Ballington, 1986). Blueberry has rapidly emerged as a major berry crop, with the United States ranking among the top three producing nations of this commodity and the leading consumer of this fruit (IBO, 2023). *V. corymbosum* is the most important species that compose the genetic background of cultivated highbush blueberries, however, this crop possesses a broad gene pool derived from wild species used during the domestication process and crop improvement (Lyrene and Ballington, 1986; Edger et al., 2022).

In this context, for a typical cultivated northern highbush blueberry (NHB) it is expected that its genetic background is comprised of approximately 90% of *V. corymbosum* (Edger et al., 2022). NHB cultivars have higher chilling requirements for flower bud break and are therefore better adapted to temperate climates where the temperature conditions allow a normal reproductive cycle. On the other hand, it is estimated that only about 70% of *V. corymbosum* compose the genetic background of southern highbush blueberry (SHB) (Brevis et al., 2008). The reduction in genomic contribution from *V. corymbosum* in SHB varieties is due to the incorporation of other *Vaccinium* species, such as *V. virgatum* and *V. darrowii*. These introgressions confer novel adaptability traits, such as reduced chilling requirements, and allow the blueberries to be grown in subtropical climates (Brevis et al., 2008; Cui et al., 2022). While the *Vaccinium* genus is highly diverse, with more than 400 species, most of the breeding efforts and genomic research to date have focused on only a handful of these species, which are considered the primary and secondary gene pools for blueberry breeding.

The successful hybridization history of blueberry involved mostly species within the *Cyanococcus* section of the *Vaccinium* genus. A modern breeding effort now is focused on increasing genetic diversity by intersectional crosses between elite blueberry cultivars and previously unused *Vaccinium* wild species. These intersectional hybrids are focused on the introgression of a new gene pool to blueberry cultivars, thereby introducing new traits, such as enhanced health-promoting benefits, tolerance to abiotic stresses, and improved flowering-related traits (Lyrene, 2011, 2016, 2018).

Among the species of interest for blueberry improvement, there is *Vaccinium stamineum* L. (2x=2n=24) (Coville, 1927), commonly known as deerberry, is the only species belonging to the section *Polycodium* within the *Vaccinium* genus. *V. stamineum* is a highly polymorphic species (Ford, 1995). This species is native to North America, and it has a wide range of habitats ranging east to west from central Florida to east Texas, and to the north up to southern Ontario, Canada (Hill, 2002; Lyrene, 2021). *V. stamineum* has similar adaptability traits compared to SHB, such as low chilling requirements, adaptation to well-drained sandy soils with low pH. In addition, a mature *V. stamineum* plant can tolerate drought stress of annual rain of 30 inches (762mm) a year, and it is also considered to be fire-tolerant and, therefore is often found in burned sites. *V. stamineum* is a perennial shrub ranging from 0.3 to 3.0 m in height, with an open canopy (Hill, 2002).

This species exhibits several intriguing traits from a plant breeding perspective that could be introgressed into elite blueberry germplasm. Some of these notable characteristics include open-flower morphology that could facilitate honey-bee pollination, natural occurrence of high soluble solids content in berries, distinctive flavor profile, and shifted harvesting season (Ballington et al., 1984; Ford, 1995; Horvat et al., 1996; Hill, 2002). However, the most remarkable trait present in some *V. stamineum* accessions is their ability to accumulate anthocyanins in the fruit pulp, which sets them apart from all other *Vaccinium* species utilized in blueberry breeding (Hill, 2002; Edger et al., 2022). This anthocyanin accumulation results in a red to deep purple internal berry color, a trait not observed in any commercial American blueberry cultivars. The goal of this study was to generate a genomic framework to facilitate research on *V. stamineum* and better understand some of the traits of interest present on this species that could be integrated into blueberry breeding.

## Material and Methods

### Plant Material

The *V. stamineum* accession named ‘AP3’ grown in 40 liters pot at the University of Florida Main Campus located in Gainesville-FL, USA. For DNA sequencing, new vegetative growth of ‘AP3’ genotype was etiolated in dark for 3 to 5 days, to increase the nuclear:plastid DNA ratio and reduce secondary metabolites, optimizing the results for the DNA extraction. Upon collection, the leaves were immediately flash frozen in liquid nitrogen, and stored in -80°C until processed. For full-length transcript isoforms sequencing (Iso-seq), non-etiolated leaves from new growth were collected, following the same collection procedures and storage. Finally, for short-reads RNA-seq, berries from six progeny accessions of *V. stamineum* ‘AP3’ were harvested at green, pink and ripe stages. Then, pulp and skin of these berries were separated and individually processed as two separate samples. The berry samples were flash frozen immediately upon collection and remained frozen at -80°C until processed.

### DNA/RNA Extraction and Sequencing

DNA extraction and PacBio HiFi sequencing was performed by the University of Wisconsin Biotechnology Center DNA Sequencing Facility (Madison, WI). First, high molecular weight DNA extraction was carried out using etiolated leaves as starting material. DNA concentration was assessed using a Quantus™ Fluorometer, while molecule sizing was verified with the Femto Pulse. The PacBio HiFi library preparation and sequencing were carried out in one SMRT cell on a PacBio Sequel IIe platform. A total of 1,167,256 HiFi reads with average length of 8741.22 bp were obtained, spanning 10,203,245,339 bp, and achieving a depth of sequencing of 17X considering the haploid genome size of 600 Mb of blueberry species. For Omni-C chromatin conformation capture, etiolated leaf samples were shipped to Cantata Bio LLC (California, USA), and the Omni-C proximity ligation library was sequenced with Illumina 150 paired-end reads. A total of 100,607,222 and 101,182,590 Omni-C paired-end reads were used to scaffold the primary and secondary haplotypes, respectively.

For the transcriptomics dataset, total RNA from both leaf and berry samples, including pulp and skin, were extracted using the Total RNA Purification Kit from Norgen Biotek Corp. (Thorold, ON, Canada), including an on-column DNAse treatment. The RNA extract quality control included NanoDrop quantification and quality assessment, fluorescent quantification with QUBIT, RNA integrity evaluation using an Agilent 2100 Bioanalyzer, and electrophoresis in a 1% agarose gel. The RNA-seq samples were submitted to Illumina TruSeq Stranded mRNA Prep Kit (Poly A selection method) and sequenced at a depth of 30M reads per sample in a NovaSeq X Series 10B (2×150). For Iso-seq, the RNA sample was submitted for the PacBio SMRTbell Library Construction, followed by sequencing in one-third of a SMRT cell on a PacBio Sequel IIe. Both RNA-seq and Iso-seq were conducted at the UF Interdisciplinary Center for Biotechnology Research (Gainesville, FL). A total of 1,494,313 Iso-Seq reads with an average size of 1,748 bp representing full length transcripts were obtained.

### Genome Assembly

First, the HiFi reads, generated from the Circular Consensus Reads (CCS), were converted to fastq format using the function *bam2fastq* loaded within the software *pacbio*/11.0.0. The initial draft assembly was performed using *hifiasm*/0.16.1 (Cheng et al., 2021). This analysis resulted in two haplotype-phased assemblies, the primary (H1) and secondary (H2) haplotypes. Then, chromatin conformation capture Omni-C reads were used for scaffolding the genomes into chromosome-size pseudomolecules using the Cantata Bio HiRise® pipeline (Putnam et al., 2016). To evaluate their synteny and identify potential misassembles, the resulting two *V. stamineum* haplotype assemblies were mapped to each other, and compared to the blueberry reference genome of the *V. corymbosum cv.* ‘Draper’ (Colle et al., 2019) using *minimap2 /2.12* (Li, 2018). The chromosome-level pseudomolecules were manually inspected for the presence of telomeric regions containing the sequences “CCCTAAA/TTTAGGG”, typical in plant genomes (Tao et al., 2024). The genome depth per haplotype was assessed by remapping the PacBio HiFi reads to the haplotype assemblies using *pbmm2*/1.13, and average coverage was estimate for each pseudomolecule individually. The 12 pseudomolecules from *V. stamineum* were renamed and reordered according to the first haplotype of the *V. corymbosum* ‘Draper’ reference genome (Colle et al., 2019). The remaining unplaced contigs were renamed and ordered by size.

### Chloroplast Sequences

To recover the scaffolds that correspond to the chloroplast sequences, the curated plastome of *V. corymbosum* SHB ‘Arcadia’ was retrieved from Fahrenkrog et al. (2022). This sequence was mapped to the *V. stamineum* assemblies using *minimap*/2.12.

### Repetitive Element Annotation

The repetitive sequences within the *V. stamineum* genome were annotated using the EDTA v.1.9.7 pipeline (Ou et al., 2019). The analysis was carried out using the default parameter except for the use of *V. caesariense* ‘W8520’ protein coding sequence (Mengist et al., 2022) as evidence to help with the annotation of transposable elements (TEs), and the “*-sensitive* 1” option that enables REPEATMODELER v.2.0 to identify any remaining TEs (Flynn et al., 2020). Additionally, the genome quality in terms of contiguity was assessed using the LTR Assembly Index (LAI) score, from the outputs of the EDTA pipeline. Once identified, the GFF file containing the positions of the repetitive sequences were softmasked using *bedtools*/2.30.0 (Quinlan and Hall, 2010).

### Protein Coding Gene Prediction and Functional Annotation

Two different approaches were used to annotate the genome: *ab initio* annotation with the web-based software helixer (https://www.plabipd.de/helixer_main.html) (Stiehler et al., 2020; Holst et al., 2023) and an evidence-based annotation using a customized *MAKER* pipeline for *V. stamineum*. The *ab initio* approach directly utilized the genome FASTA file to predict gene models based on pre-computed algorithms from other plant species.

For the homology-based approach, it was necessary to assemble a transcriptome for *V. stamineum* that was used as evidence to train and predict gene model calls. First, RNA-seq reads from berry tissues were mapped to each haplotype of the *V. stamineum* genome using *hisat2*/ 2.2.1 (Kim et al., 2019) and generated BAM files with *samtools*/1.19.2 (Danecek et al., 2021). Then, we used *stringtie*/2.2.3 (Kovaka et al., 2019) with the *–mix* flag to combine both, RNA-seq reads from berries and the Iso-seq reads from leaves and generate a GTF file with the transcript coordinates in the reference genome. This GTF file was converted to GFF format with *gffread*/0.12.7 (Pertea and Pertea, 2020) and later to a FASTA file containing each transcript sequences using *bedtools*/2.30.0 (Quinlan and Hall, 2010).

The soft-masked genome from the EDTA pipeline was then used as input for the annotation of protein coding genes. The transcriptomic datasets from *V. stamineum* and homologous protein sequences were used as evidence in the *MAKER* pipeline (Cantarel et al., 2008; Badalà et al., 2014). Protein evidence from both haplotypes of *V. caesariense* ‘W8520’ genome were included to support the training of gene model prediction software. This approach should provide evidence of gene models that were not captured in the *V. stamineum* transcriptome assembly previously described. The gene prediction was carried out for each *V. stamineum* haplotypes independently. In the first round of gene predictions by *MAKER*, both, the *V. stamineum* transcriptome and proteome from *V. caesariense* were used as evidence to perform BLAST searches and *EXONERATE* (Slater and Birney, 2005) annotated initial homology-based predictions. From here, Hidden Markov Models were trained for *SNAP* (Korf, 2004) and *Augustus* (Stanke et al., 2008) based on the predictions performed in the previous annotation iteration. Two more rounds of *ab initio* gene model predictions were performed.

The gene annotation was polished by using a dataset containing reviewed and curated genes for the *Viridiplantae* clade was downloaded from UniProt (www.uniprot.org/, access on: September 19^th^ 2024) containing 41,742 entries. We used the *maker-functional* scripts to identify conserved domains and perform BLAST searches that provided support for the gene models predicted by the MAKER pipeline. Functional annotation was also carried out using PANNZER2 (Törönen et al., 2018) and eggNOG mapper (Huerta-Cepas et al., 2019) to retrieve Gene Ontology (GO) terms, Kyoto Encyclopedia of Genes and Genomes (KEGG) and Functional Clusters of Orthologous Groups (COG).

To generate a set a of high confidence (HC) genes, the gene models called by MAKER were filtered based on the presence of conserved domains, and by a homology-based search against the UniProt *Viridiplantae* database using BLASTP with an e-value cutoff of 10^−6^. In addition, this gene set was further filtered for models with an Assembly Evaluation Distance (AED) score ≤ 0.1, meaning these predictions had evidence to support the annotation and did not rely solely in *ab initio* prediction.

### Assessment of Genome and Transcriptome Resources

The newly generated genome assemblies and gene annotations for two haplotypes were evaluated to ensure their quality. BUSCO (Benchmarking Universal Single-Copy Orthologs) (Manni et al., 2021b, 2021a) analysis was conducted on both the genome and transcriptome level to assess the overall completeness of the genic content in each assembly, based on the *embryophyta_odb10* dataset for land plants, containing 1614 single-copy orthologs. This approach allowed for the evaluation of gene fragmentation or missing information present in the assembled genomes and the gene annotation quality reflected in their corresponding transcriptome. Additionally, the intergenic regions were assessed using the LAI (LTR Assembly Index) score (Ou et al., 2018), which measures the abundance and fragmentation level of LTR retrotransposons, which are abundant transposable elements in plants.

## Results

### Genome Assembly

The PacBio HiFi long reads were used to generate a draft genome assembly that was partially haplotype-phased as switches between haplotypes are expected based on the amount of sequencing information (Sharma et al., 2022). The assembled genomes had a N50 of 0.549 Mb and 0.569 Mb, for primary (H1) and secondary (H2) haplotypes, respectively. In addition, the haplotype H1 spawned 3,427 contigs and had the longest contig of 4.59 Mb in length, while the haplotype H2 spawned 1,899 contigs and had the longest contig of 5.77 Mb in length (Table 1). The draft assemblies were then submitted to scaffolding using Omni-C reads that uses chromatin ligation distances to accurately place contigs into chromosome-scale scaffolds. This step considerably improved contiguity. After scaffolding, the primary haplotype spawned 529.16 Mb and 2,044 scaffolds, while the secondary haplotype spawned 493.82 Mb over 557 scaffolds. Although, the number of scaffolds remained large, the 12 largest scaffolds for both haplotypes were assembled to chromosome level pseudomolecules corresponding to the 12 chromosomes in *Vaccinium* spp. (Table 1). For the first haplotype, the first 12 pseudomolecules correspond to 89.04 % of the total assembly size, while for the second haplotype 94.28% of the assembled genome was comprised within the pseudochromosomes. Finally, the 12 pseudochromosomes of both haplotypes were ordered based on the blueberry reference genome cv. ‘Draper’ from Colle et al. (2019). The remaining contigs were ordered by size from largest to smallest.

**Table 1.** Assembly statistics the final genome version, before and after scaffolding using Omni-C chromatin conformation data.

| Parameters | Primary Haplotype (H1) | Secondary Haplotype (H2) |
| --- | --- | --- |
| *Before Scaffolding* |  |  |
| # Contigs/Scaffolds | 3427 | 1899 |
| Largest Contig (Mbp) | 4.59 | 5.77 |
| N50 (Mbp) | 0.549 | 0.569 |
| *After Scaffolding* |  |  |
| # Contigs/Scaffolds | 2044 | 557 |
| Total Length (Mbp) | 529.16 | 493.82 |
| Largest Contig (Mbp) | 47.3 | 46.45 |
| GC (%) | 39.2 | 38.49 |
| N50 (Mbp) | 38.82 | 38.8 |
| L50 | 7 | 6 |
| BUSCO | 97.4% [ S:91.4%, D: 6.0%] | 96.7% [ S:91.0%, D: 5.7%] |
| LAI | 19.63 | 23.51 |
| Number of N's per 100kb | 26.29 | 27.32 |
| Annotated genes | 62373 | 58553 |
| Telomere-to-telomere assembly (Chrom) | 1, 2, 7 ,11 and 12 | 2, 8, 9 and 11 |
| Telomere at the start (Chrom) | 4, 5, 6, 9 and 10 | 1, 3, 4, 5, 7 and 12 |
| Telomere at the end (Chrom) | 3 and 6 | 6 and 10 |

Additionally, after scaffolding, the 13^th^ largest scaffold (*VstamH1_sc13* and *VstamH2_sc13*) had a significant size drop, with a reduction of more than 30Mb for both haplotypes. By aligning this 13^th^ largest scaffold to the first 12 pseudochromosomes in each haplotype and by visual inspection of the sequences, it is possible to observe that these scaffolds are comprised of several repetitive sequences that were compiled together during the assembly process, supported by multiple alignments with several loci in the 12 pseudochromosomes.

The presence of telomeric regions, characterized by the canonical ‘CCCTAAA/TTTAGGG’ repeats in plants, were visually inspected in the Integrated Genome Browser (IGV) software (Table 1). Interestingly, several chromosomes were assembled at a telomere-to-telomere resolution, including five pseudomolecules in the primary haplotype and four pseudomolecules in the secondary haplotype. All remaining chromosomes in each assembly had telomeric sequences identified in either the start or the end of the pseudomolecules.

Chloroplast sequences were identified by mapping the chloroplast assembly of SHB ‘Arcadia’ chloroplast from Fahrenkrog et. al (2022) to each haplotype of *V. stamineum*. The sequence alignment showed that the scaffold ‘*VstamH1_sc34*’ and ‘*VstamH2_sc40*’, both correspond to chloroplast sequences of *V. stamineum*, spawning exactly 162,154bp in both haplotypes. This analysis showed that we were able to recover the entire chloroplast genome within only one scaffold.

In terms of base calling, each scaffold was assessed independently within each haplotype. Overall, high scores of quality value (QV) were achieved, suggesting that the assembly should have minimal errors. For reference, a QV > 40 corresponds to a probability of error of 1 in 40 billion. In the primary haplotype, chromosome 1 had the lowest QV score of 41.62, while the chromosome 5 had the highest QV score of 58.05. In case of the secondary haplotype, all QV scores were above 55, demonstrating the high confidence in base calling for both haplotypes. Finally, the PacBio Reads were remapped to the assemblies, showing that the pseudomolecules were assembled with similar sequencing depth. For haplotype H1, chromosome 11 had the lowest overall depth of coverage while chromosome 05 had the highest overall coverage, 16.34 and 17.17X, respectively. For haplotype 2, the chromosome with the lowest depth was 06, with 16.54X while chromosome 05 also showed the highest depth with 17.22X.

The BUSCO analyses revealed that our newly assembled genome was fairly complete reaching a BUSCO of 98.9% at the genome level assessment, with the BUSCO of the primary haplotype C:97.4%[S:91.4%,D:6.0%] and the secondary haplotype C:96.7%[S:91.0%,D:5.7%] (Figure 1).

**Figure 1.**
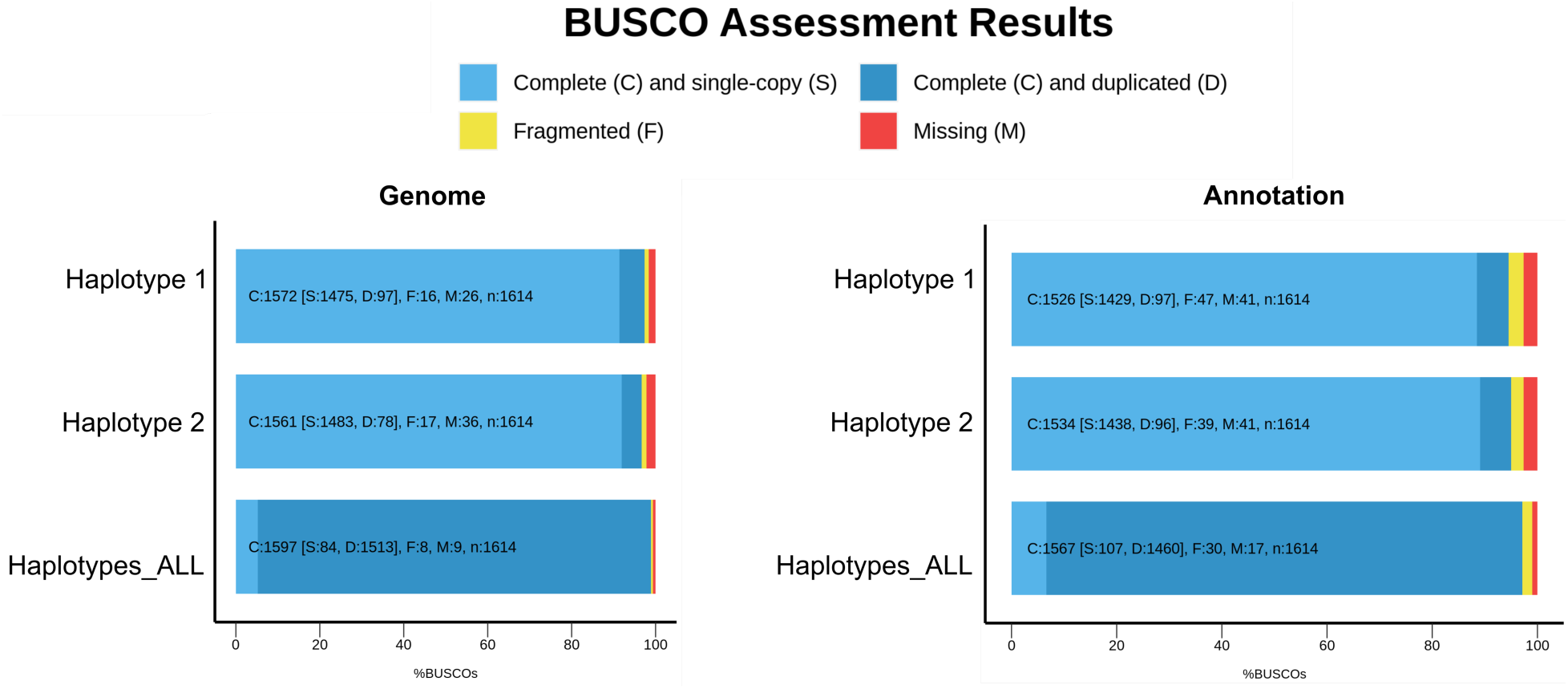
Assessment of BUSCO scores for each *V. stamineum* haplotype and their combination (Haplotypes_ALL) at the genome and annotation level.

### Transposable Element and Protein Coding Gene Annotation

TEs were identified using the EDTA pipeline individually for each haplotype. For this step, we included protein evidence from *V. caesariense* to support the analysis and mitigate TEs being overly annotated. A total of 39.77% and 42.38% of the sequences from haplotype H1 and haplotype H2 were identified as repetitive elements, respectively (Table 2). The most abundant types of repeats were LTR-Gypsy, nonTIR-helitron and TIR-Tc1_Mariner corresponding to 7.65%, 4.57% and 4.53%, respectively, in the primary haplotype. The secondary haplotype exhibited some differences in the abundance of transposable elements. LTR-Gypsy remained the most common, followed by TIR-Mutator, while LTR-Copia and TIR-Tc1_Mariner contributed equally to the number of repetitive sequences, each accounting for 8.21%, 4.74%, 4.69%, and 4.69%, respectively. Finally, LAI score was calculated as an assessment of completeness of intergenic regions, resulting in a score of 19.63 for the primary haplotype, just short of being considered a gold standard assembly, and 23.51 for the secondary haplotype, therefore characterizing it as gold standard genomes for this parameter (Table 2). The repetitive sequences identified were used to generate a softmasked version of the genome, as an input for protein coding gene annotation.

**Table 2.**
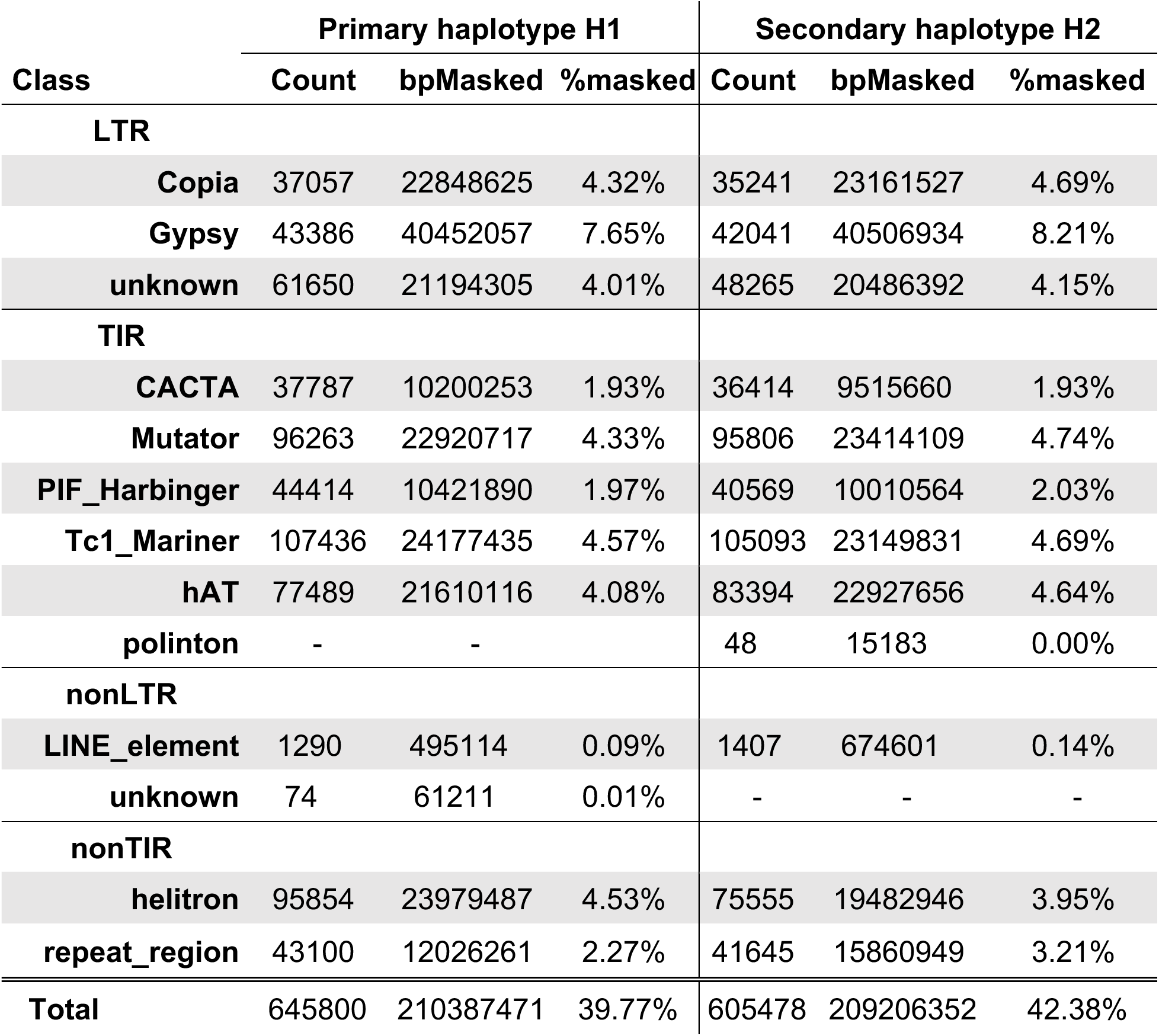
Repetitive sequences identified in the V. stamineum haplotypes using the EDTA pipeline.

| Class | Primary haplotype H1 |  |  | Secondary haplotype H2 |  |  |
| --- | --- | --- | --- | --- | --- | --- |
|  | Count | bpMasked | %masked | Count | bpMasked | %masked |
| **LTR** |  |  |  |  |  |  |
| **Copia** | 37057 | 22848625 | 4.32% | 35241 | 23161527 | 4.69% |
| **Gypsy** | 43386 | 40452057 | 7.65% | 42041 | 40506934 | 8.21% |
| **unknown** | 61650 | 21194305 | 4.01% | 48265 | 20486392 | 4.15% |
| **TIR** |  |  |  |  |  |  |
| **CACTA** | 37787 | 10200253 | 1.93% | 36414 | 9515660 | 1.93% |
| **Mutator** | 96263 | 22920717 | 4.33% | 95806 | 23414109 | 4.74% |
| **PIF_Harbinger** | 44414 | 10421890 | 1.97% | 40569 | 10010564 | 2.03% |
| **Tc1_Mariner** | 107436 | 24177435 | 4.57% | 105093 | 23149831 | 4.69% |
| **hAT** | 77489 | 21610116 | 4.08% | 83394 | 22927656 | 4.64% |
| **polinton** | - | - |  | 48 | 15183 | 0.00% |
| **nonLTR** |  |  |  |  |  |  |
| **LINE_element** | 1290 | 495114 | 0.09% | 1407 | 674601 | 0.14% |
| **unknown** | 74 | 61211 | 0.01% | - | - | - |
| **nonTIR** |  |  |  |  |  |  |
| **helitron** | 95854 | 23979487 | 4.53% | 75555 | 19482946 | 3.95% |
| **repeat_region** | 43100 | 12026261 | 2.27% | 41645 | 15860949 | 3.21% |
| **Total** | 645800 | 210387471 | 39.77% | 605478 | 209206352 | 42.38% |

The gene annotations were generated using two software Helixer and MAKER. Helixer is an *ab initio* prediction software, it does not require evidence to support gene annotation, and it provides a full annotation within a few hours. The Helixer pipeline for the primary haplotype predicted 35,188 gene models with an transcriptome level BUSCO of C:92.9%[S:87.8%,D:5.1%]. Conversely, the MAKER pipeline is benefited from protein and transcript evidence combining both, homology-based and *ab initio* predictions, but it takes several days and multiple iterations to achieve a final output. Transcript evidence was obtained from short reads Illumina RNA-seq from berries and long-reads PacBio Iso-seq from leaves of *V. stamineum*. These datasets were combined and used to assemble a transcriptome as evidence to support gene prediction. From this approach, a total of 68,437 transcripts were assembled. Finally, by combining the newly assembled *V. stamineum* transcriptome and protein evidence from *V. caesariense*, a total of 62,397 and 58,553 gene models were predicted for haplotype H1 and haplotype H2, respectively, over three rounds of annotation using the *MAKER* pipeline, retraining the HMM of *Augustus* and *SNAP* between iterations (Table 1). Additionally, to the increase in gene model predictions, the BUSCO scores for both haplotypes also increased in the *MAKER* annotations, corresponding to C:94.5%[S:88.5%,D:6.0%] and C:95.0%[S:89.1%,D:5.9%] for the primary and secondary haplotypes, respectively (Figure 1).

Finally, from the functional annotation using eggNOG mapper, it was possible to identify over 40,000 genes with assigned COG categories of which approximately 11,000 were classified with unknown function (Figure 2).

**Figure 2.**
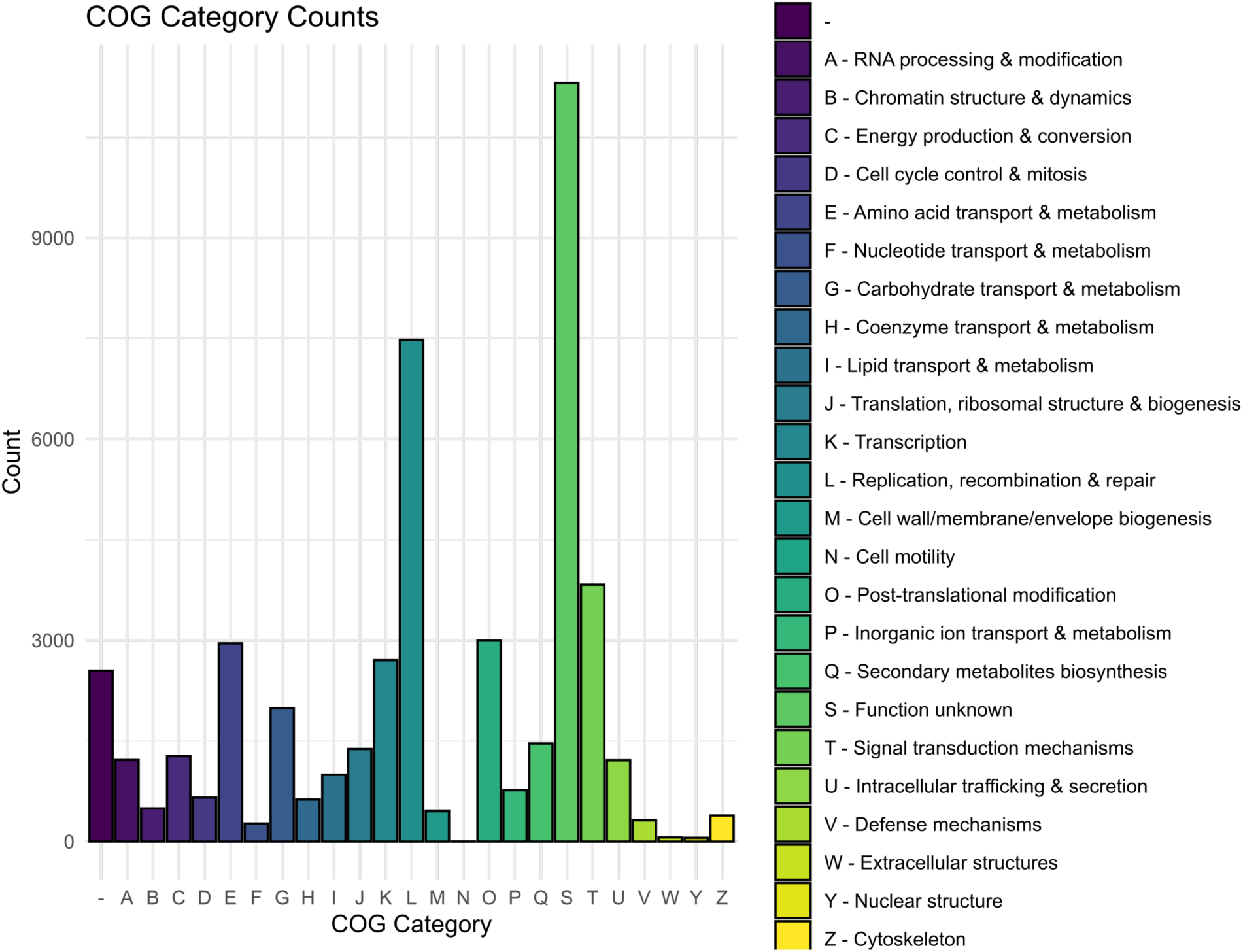
Cluster of Orthologs (COGs) counting per category based on the functional annotation given by EGGNOG mapper in *V. stamineum* primary haplotype.

The subset of HC genes reflect genes models that were called based on the presence of transcript evidence, conserved domains and homology detection. In the primary haplotype, the HC set was composed of 23,898 genes models. This set of genes can be taken as a reliable set for further functional and evolutionary analyses.

### Synteny Between V. stamineum and V. corymbosum

Nucleotide alignments between the two haplotypes of *V. stamineum* showed that they were highly collinear and syntenic to each other as expected. However, the end of chromosome 1 in the haplotype H1 aligned to the beginning of the chromosome 1 in haplotype H2, due to either a translocation that caused such rearrangement or a possible misassembly (Figure 3A).

**Figure 3.**
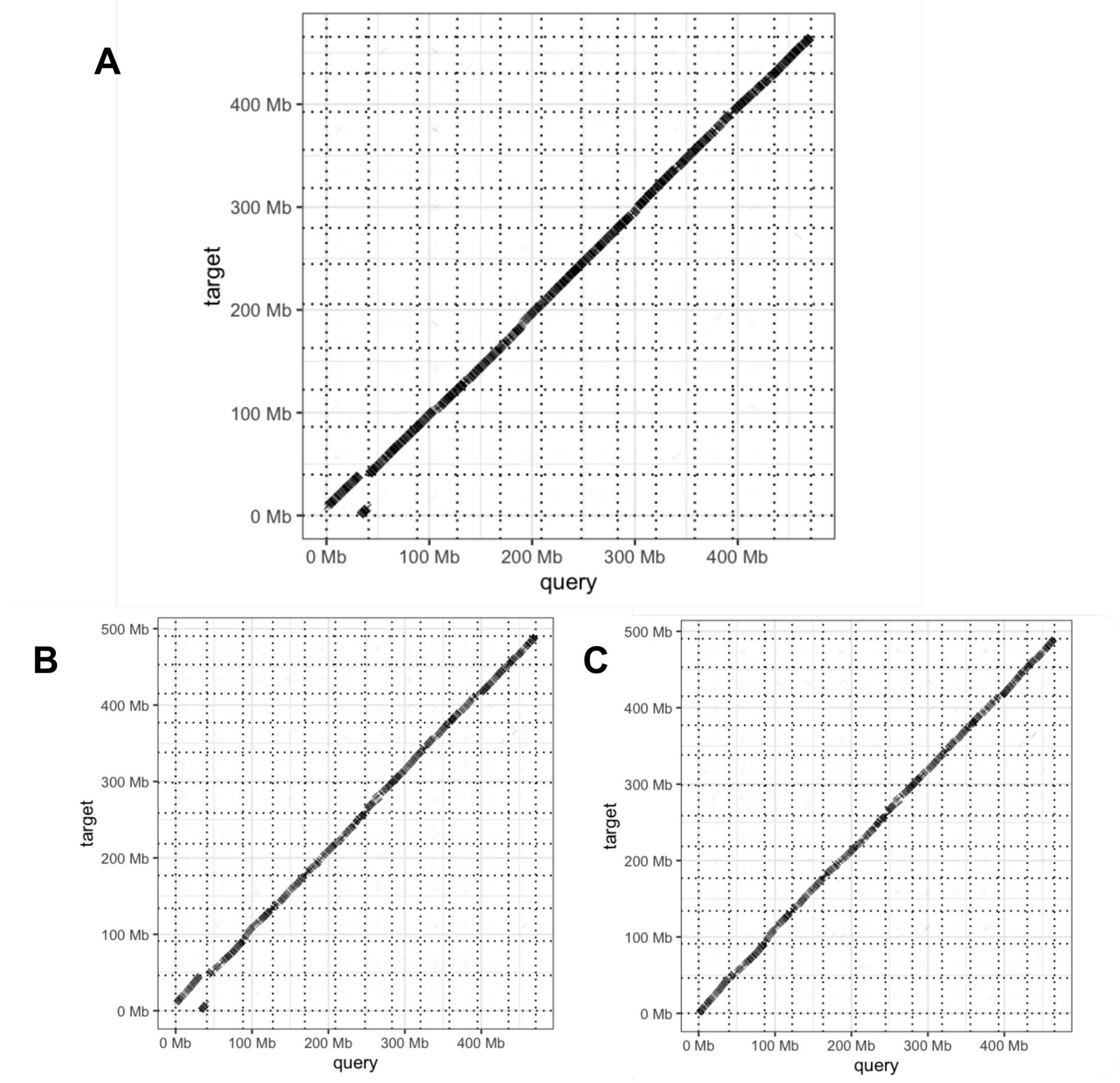
Collinearity between the two haplotypes of the *V. stamineum* assembly and *V. corymbosum* cv. ‘Draper’ using nucleotide alignments (>5 kb) generated with *minimap2*. A) Collinearity between the two haplotypes of V. stamineum, with as haplotype 1 as query (X axis) and haplotype 2 (Y axis). B) Collinearity between haplotype 1 of *V.* stamineum (X axis) and haplotype 1 of *V. corymbosum* cv. ‘Draper’ (Y axis). C) Collinearity between haplotype 2 of *V. stamineum* (X axis) and haplotype 1 of *V. corymbosum* cv. ‘Draper’ (Y axis).

As previously mentioned, the 12 chromosome-level pseudomolecules carry most of the genetic information in each genome, and both haplotypes seem to carry the same information throughout the genome, except for a few loci where primary haplotype seems to be more completed compared to its counterpart (Figure 4).

**Figure 4.**
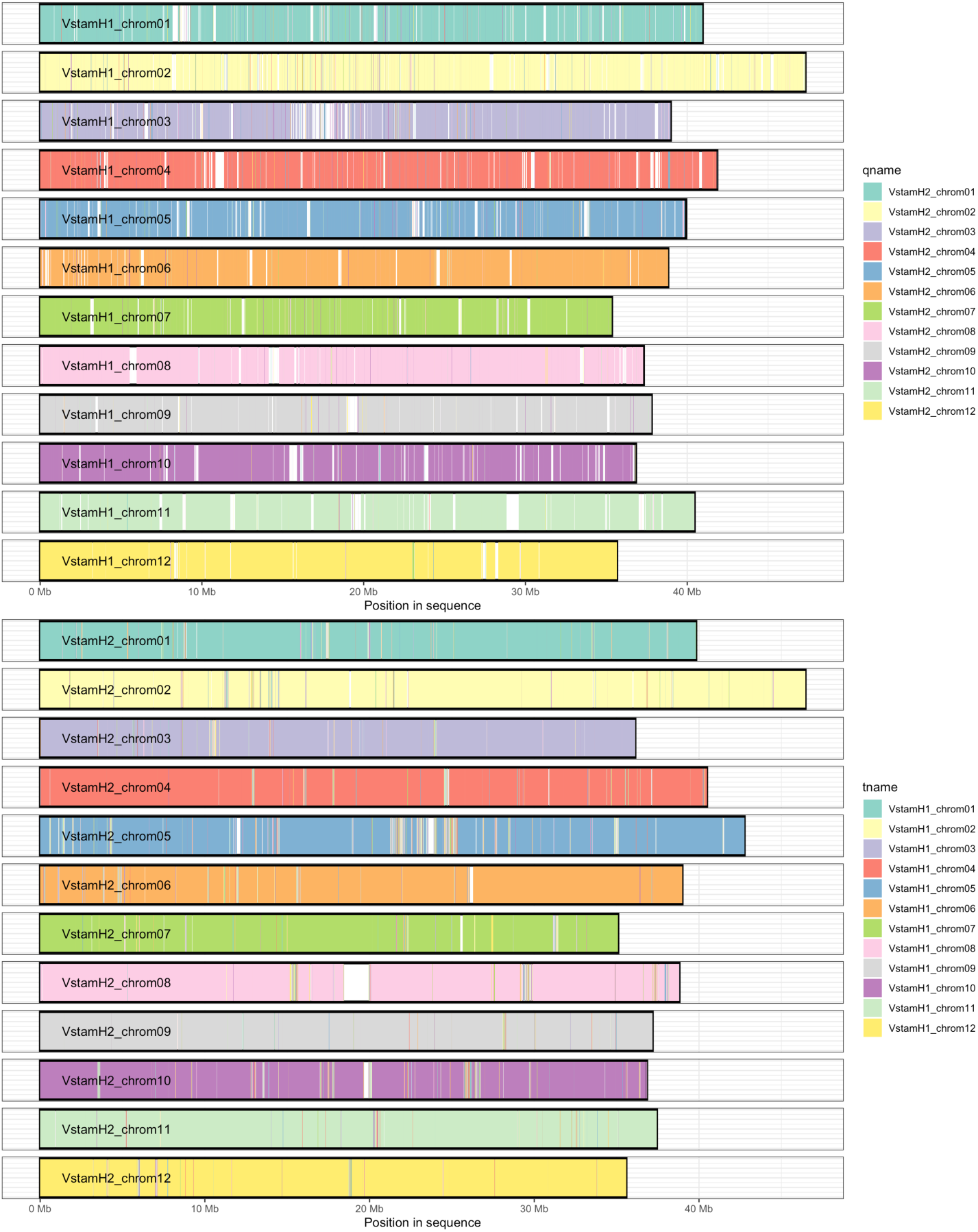
Synteny and haplotype completeness. On top, the chromosomes from the primary haplotype H1, are represented by boxes displaying their sizes in Mb. The color inside of each chromosome box is given by nucleotide alignments of >5 kb with the secondary haplotype H2. White space are regions present in H1 that do not possess a correspondent alignment with H2. On bottom, the chromosomes from the secondary haplotype H2, are represented by boxes displaying their sizes in Mb. The color inside of each chromosome box is given by alignments of >5 kb with the primary haplotype H1. White space are regions present in H2 that do not possess a correspondent alignment with H1.

Finally, nucleotide alignment showed that both haplotypes of *V. stamineum* were also highly collinear and syntenic to the first haplotype of NHB ‘Draper’ genome. The putative rearrangement in chromosome 1 of the primary haplotype persisted in this comparison. This result suggests a possible structural variation or assembly artifact and further analyses must be performed to clarify the nature of such discrepancies (Figure 3 B and C).

### Discussion Generation of Genomic Resources for *Vaccinium* Species

The *Vaccinium* genus is very large, and its species richness represents a reservoir of genetic diversity available for improving blueberry and cranberry crops. However, in the past decades, the breeding efforts in *Vaccinium* crops have focused on outsourcing genetic material from only a few wild relatives, resulting in an underexplored range of wild species that could bring novel traits to cultivar improvement (Edger et al., 2022). In this context, generating more genomic resources for wild crop relatives is critical for gene and novel trait discovery and accelerate breeding endeavors. Therefore, the goal of this study was to generate a genomic framework for *V. stamineum*, which is a promising species for *de novo* domestication and introgression into blueberry.

In this study, we generated the first haplotype-phased reference genome of a *V. stamineum* accession called ‘AP3’. This genotype is especially interesting due to its ability to accumulate anthocyanin in the fruit pulp, a trait that is currently not observed in any American blueberry cultivar. For the assembly of the genomes, in normal conditions, the amount of data generated by one SMRT cell should fall between 12 to 19 Gb of CCS, resulting in 20 to 30X coverage of the genome. This depth should be enough to characterize a diploid genome (Motazedi et al., 2018). The genome size of the *V. stamineum* reference genome here assembled aligned with the range reported by Redpath et al. (2022) for this species. In agreement with this, the final assemblies for the primary and secondary haplotypes yielded a similar genome size, with most of the genome comprised within 12 pseudomolecules corresponding to the 12 base chromosomes present in *Vaccinium* species. The detection of telomere repeat sequences at the termini of most the pseudomolecules also indicates we were close to a telomere-to-telomere (T2T) sequence assembly.

Both haplotypes assembled for *V. stamineum* were highly syntenic to each other, with the primary assembly encompassing a more complete representation (less gaps per total length) of a haploid *V. stamineum* genome than the secondary haplotype. However, the primary haplotype of chromosome 1 exhibited a translocation at the end of the chromosome when aligned to the secondary haplotype. The same pattern was observed when the primary haplotype was aligned against the primary haplotype of the *V. corymbosum* cv. ‘Draper’ assembly. Whether the observed translocation is a genuine feature of *V. stamineum* haplotype 1 or an artifact of the assembly process remains uncertain, and further validation is required.

The *V. stamineum* genome exhibited repetitive sequences comprising 39.77% and 42.38% of the primary and secondary haplotypes, respectively. This is slightly smaller when compared to the repetitive content reported in other *Vaccinium* species, including *V. corymbosum* (44.3%) (Colle et al., 2019), *V. caesariense* (45.3%) (Mengist et al., 2022), *V. darrowii* (45.74%) (Yu et al., 2021), *V. myrtillus* (47.46%) (Wu et al., 2021), and *V. macrocarpon* (50.49%) (Kawash et al., 2022). It is noteworthy that the genome size of *V. stamineum* was predicted to be approximately 50 Mb smaller than most *Vaccinium* genomes, and interestingly, the total amount of repetitive elements was reduced in this species. In contrast, the number of protein-coding genes predicted in the two *V. stamineum* haplotypes was higher than the estimates in other *Vaccinium* species, indicating that the smaller genome size was not accompanied by a reduction in gene content. For instance, the *V. corymbosum* genome was estimated to have 128,559 total protein-coding genes for the tetraploid assembly, with approximately 32,140 genes per haplotype, while the haploid representation of *V. stamineum* had 62,397 predicted gene models. Genome sizes are known to vary within species and in comparison with crop wild relatives (Kreiner and Wright, 2018). Among the main reasons why genome sizes fluctuations happens there are: variation the TE composition, polyploidization and diploidization events, and accumulation of mutations such as large indels (Grover et al., 2004; Kreiner and Wright, 2018). On the other hand, variation in the gene content could be strictly related to diversification in adaptability traits to different environments but also just an artifact of differing gene annotation strategies, including annotation software as well as provided evidence to support the annotation (Nachtweide et al., 2024).

Although there were significant differences found in the gene content of *V. stamineum*, the BUSCO analysis revealed that our newly assembled genome was fairly complete reaching a BUSCO of 98.9% at the genome level assessment. Similar BUSCO scores have been reported for other *Vaccinium* spp., such as *V. darrowii* (from 88.9% to 93.49%) (Yu et al., 2021; Cui et al., 2022); *V. myrtillus* (96.6%) (Wu et al., 2021), *V. macrocarpon* (93.4%) (Kawash et al., 2022), *V. corymbosum* (97%)(Colle et al., 2019) and *V. caesariense* (>99%) (Mengist et al., 2022). It is noteworthy that the BUSCO scores were not assessed using the same database, which may mislead the comparison of the gene content completeness among these genomes. Our results in *V. stamineum* were performed using the most up to date database for land plants, ‘*embryophyta_odb10*’, which is the same database used in the BUSCO assessment of both *V. darrowii* genomes, and *V. myrtillus*. In terms of the transcriptome, the *MAKER* pipeline achieved a higher BUSCO score in the primary haplotype, compared to the Helixer annotation, C:94.5%[S:88.5%,D:6.0%] and C:92.9%[S:87.8%,D:5.1%], respectively. Furthermore, the *MAKER* pipeline predicted 62,373 gene models compare to 35,188 from Helixer. The variation in predicted gene models, can be explained due to the fact that *MAKER* was trained on *V. stamineum* datasets, thus potentially enhancing the sensitive of this method. Based on these results, we concluded that the evidence-based approach in gene annotation outperformed the *ab initio* model.

Altogether, our results increase the available genomic resources that can be used to understand the genetic diversity within the *Vaccinium* genus. In this study, we present the first reference genome for *V. stamineum*, the only species that composes the *Polycodium* section of the *Vaccinium* genus. Future research may use the resources we generate here to further explore the potential of this species to be introgressed into blueberry breeding or even to support the domestication of this species as a crop of its own.

## Data availability

Data is available upon request.

## Acknowledgements

This work was supported by the UF royalty fund generated by the licensing of blueberry cultivars.

